# Stimulation of Locus Coeruleus Noradrenergic System Modulates Sensory Processing and Brain State in two Different Time Scales

**DOI:** 10.1101/2020.07.09.188615

**Authors:** Zeinab Fazlali, Yadollah Ranjbar-Slamloo, Erfan Zabeh, Ehsan Arabzadeh

## Abstract

Locus Coeruleus (LC) noradrenergic system has widespread projections throughout the brain and affects sensory processing. LC modulation of sensory-evoked cortical activity and brain state is documented by electrical micro-stimulation, optogenetic experiments, and the local application of norepinephrine (NE). The temporal profile of the LC modulation of sensory response and brain state is not well characterized. Our goal in this study is to characterize this modulation. Here, we recorded neuronal activity from the barrel cortex (BC) of urethane-anesthetized rats while combining LC micro-stimulation with brief mechanical deflections of the whiskers at 10 different time lags (50-500 ms). We recorded spikes and local field potentials to quantify the neuronal activity and the brain state. LC micro-stimulation exhibited a biphasic effect on spontaneous activity of the BC: a period of suppression followed by a period of excitation. We observed a similar effect on the sensory-evoked response: at 50-ms lag, the evoked response decreased while at 150-ms lag, the early evoked response was facilitated. At 150 to 350-ms time lags, LC micro-stimulation caused a combined facilitation followed by suppression of the evoked response. In contrast to the fast transient effect of LC stimulation on BC spiking activity, brain state modulation started later and lasted longer. LC stimulation suppressed low-frequency activities that are associated with low arousal states. In summary, we found that LC modulation affects cortical processing of sensory inputs and the brain state at different time scales which are likely to involve distinct circuit mechanisms.

## Introduction

Locus Coeruleus (LC), a major neuromodulatory nucleus and the main source of norepinephrine (NE) in the brain, has widespread projections throughout the cortex and affects neuronal activity, network dynamics, and task-relevant perceptual behavior (Berridge and Waterhouse, 2003; Sara, 2009; Sara and Bouret, 2012; Waterhouse and Navarra, 2019). Previous studies have revealed a prominent role of LC-NE system in controlling behavioral and brain states (Aston-Jones and Bloom, 1981a; Aston-Jones and Cohen, 2005; Berridge and Waterhouse, 2003; Carter et al., 2010; Fazlali et al., 2016; Hayat et al., 2020) and cognitive effort (Eldar et al., 2013; Gottlieb et al., 2020; Silvetti et al., 2021, 2017) as well as modulation of sensory processing (Berridge and Waterhouse, 2003; Devilbiss and Waterhouse, 2011, 2004; Edeline et al., 2011; Fazlali and Ranjbar-Slamloo, 2021; Lecas, 2004; Manunta and Edeline, 2004; Rodenkirch et al., 2019; Waterhouse et al., 1998). Inhibition of both spontaneous and sensory-evoked cortical activity was documented in classic experiments that used either LC electrical micro-stimulation or local application of norepinephrine (Foote et al., 1975; Phillis and Kostopoulos, 1977). However, later studies reported more diverse effects on sensory response following LC electrical micro-stimulation or local NE application in the sensory cortex and thalamus (Berridge and Waterhouse, 2003; Devilbiss and Waterhouse, 2011, 2004; Edeline et al., 2011; Lecas, 2004; Manunta and Edeline, 2004; Vazey et al., 2018; Waterhouse et al., 1998; Waterhouse and Navarra, 2019). The facilitation, suppression, and the mixed effects of LC micro-stimulation on neuronal activity were found both in anesthetized (Bouret and Sara, 2004; Lecas, 2004; Manunta and Edeline, 2004) and awake rats (Devilbiss et al., 2006; Devilbiss and Waterhouse, 2011, 2004). In particular, the net effect of LC-NE activation is considered to be an enhanced “signal-to-noise ratio” of sensory responses in multiple cortical regions (Berridge and Waterhouse, 2003; Edeline, 2012; Foote et al., 1975; Hirata et al., 2006; Jiang et al., 1996; Waterhouse et al., 1998). In the olfactory bulb LC activation suppressed spontaneous activity while increasing the evoked response of the neurons (Jiang et al., 1996). In somatosensory cortical neurons, LC micro-stimulation enhanced sensory-evoked responses without a significant effect on the spontaneous firing rate (Bouret and Sara, 2002; Waterhouse et al., 1998). The maximum enhancement effect on the evoked somatosensory response was observed around 200-300 ms and diminished beyond 400-500 ms (Waterhouse et al., 1998).

Our aim here was to characterize the temporal profile of LC stimulation on neuronal spike activity in the primary sensory cortex and the consequent effects on sensory coding efficiency. In addition, we aimed to quantify the effect of micro-stimulation on cortical state changes. We, therefore, recorded spiking activity and local field potential in the rat barrel cortex and examined the LC-NE modulatory effects at several intervals between LC and sensory (whisker) stimulations.

## Material and Methods

### Surgery and electrophysiological recording

Thirty-two adult male Wistar rats, weighing 270-390 g were used in this study. All experiments were approved by the animal care and experimentation committee of the Institute for Research in Fundamental Science (IPM). Anesthesia was induced by intraperitoneal administration of urethane (1.5 g/Kg), monitored by the hind paw and corneal reflexes, and maintained with supplemental doses of urethane (0.1 g/Kg), if necessary. Two craniotomies were performed on the right hemisphere to provide access to BC (5 x 5 mm; centered at 2.6 mm posterior and 5 mm lateral to Bregma) and LC (4 x 4 mm; centered at 10.8 mm posterior to Bregma and 1.4 mm lateral to the midline). To facilitate access to LC, the rat’s head was tilted down by about 14 degrees (Bouret and Sara, 2002).

We recorded neuronal activity in BC while electrically stimulating ipsilateral LC (Figure 1A). BC neuronal activity was acquired with single tungsten microelectrodes (1-2 MΩ, FHC Inc., USA). The unit’s principal whisker was determined by manual stimulation of individual whiskers. Layer 4 of the barrel cortex was targeted based on the depth of penetration from the surface of the exposed dura. Data were collected at a sampling rate of 30 kHz and filtered on-line by applying a band-pass filter (300–6000 Hz). Spikes were sorted off-line using principal component analysis implemented in MATLAB (Math Works). In total, 49 multi-units were extracted from BC recordings.

**Figure 1.**
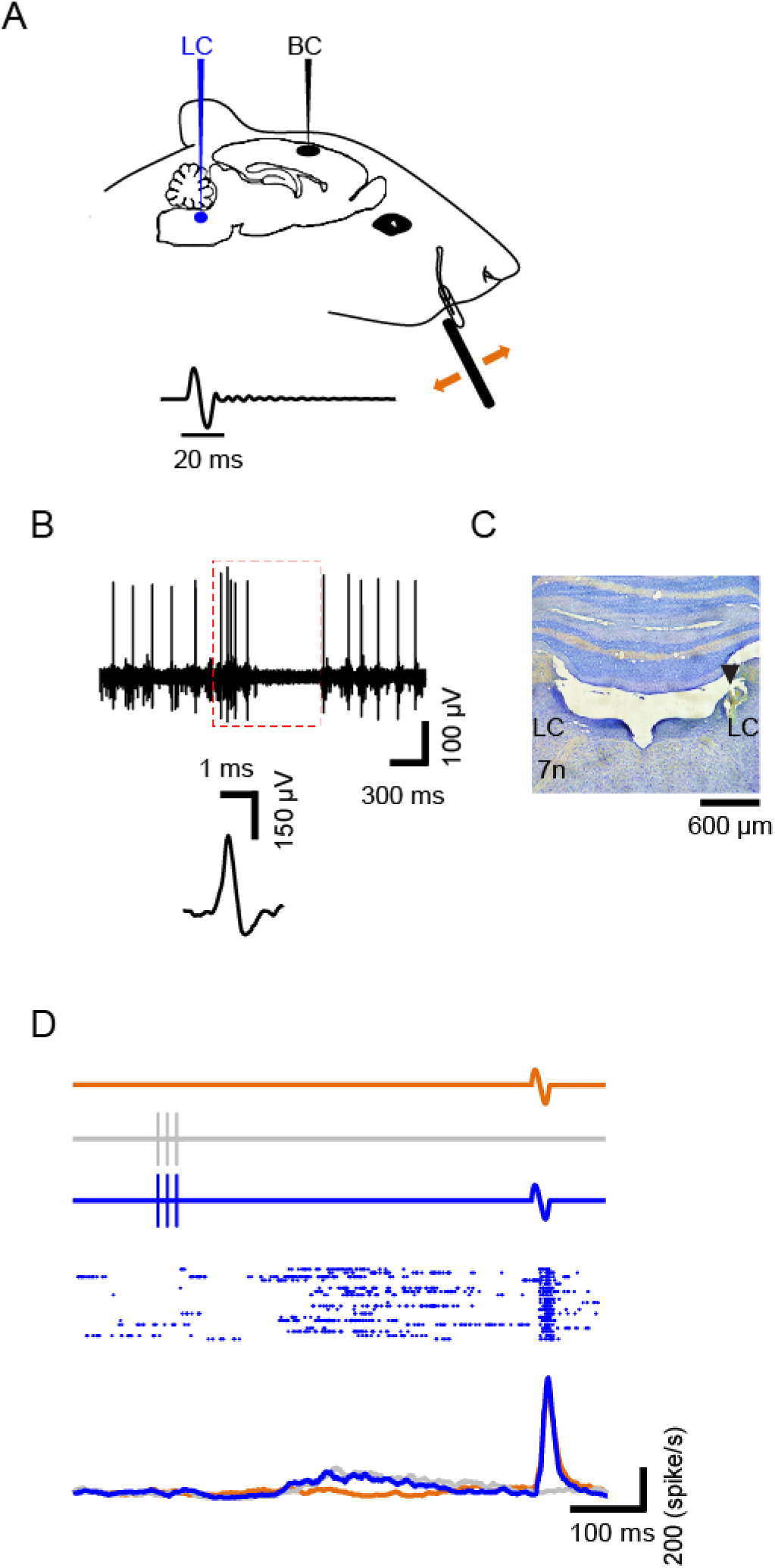
Schematic illustration of the experiment. (**A**) Neuronal activity was recorded from the Barrel Cortex (BC) while a brief mechanical vibration was applied to the contralateral whisker at various intervals relative to micro-stimulation of the Locus Coeruleus (LC). For sensory stimulation, a full-cycle sinusoidal deflection was applied to the contralateral whisker. **B** and **C** show respectively electrophysiological and histological criteria to identify LC for microstimulation (previously published under {add the license type and reference here}). (**D**) Trial types: whisker deflections but no LC micro-stimulation in brown; those with LC micro-stimulation but no whisker deflections in gray; and those that contained both LC micro-stimulation and whisker deflections in blue. Raster plot shows a typical multi-unit activity in a LC micro-stimulation-deflection trials. Lower PSTHs show the firing rate corresponding to each trial type. This color convention will be used henceforth.

### Whisker stimulation

A single cycle of 80-Hz sinusoidal waveform was delivered to the principal contra-lateral whisker at multiple amplitudes using a piezoelectric device (NikTek, Tehran, Iran). Stimuli were generated in MATLAB and presented through the analog output of the sound card at a sampling rate of 96 kHz. The principal whisker was placed in a cannula connected to the piezo with a 2 mm distance from the tip of the cannula to the base of the whisker. To ensure precise whisker stimulation, we used an infrared optic sensor to calibrate the movement of the piezo. We measured the piezo movement to confirm that it accurately followed the voltage command with minimal resonation. For the range of stimulus intensities applied here (6-60 μm), the post-deflection resonance was negligible (<6% of maximum amplitude) and was not detectable beyond 70-80 ms. To select a near-threshold range of stimuli, at the beginning of each recording session, we applied 10 levels of deflection from a relatively wide range of amplitudes (0-60 μm with 6 μm steps, 50 repetitions). A Nuka-Rushton function was fitted to the average spike count to characterize the neuronal response function. Threshold (T) was defined as the inflection point of this function – i.e. the stimulus amplitude that produced half of the maximum response (Fazlali et al., 2016).

In a subset of experiments (n=33), the threshold stimulus was applied to the whisker at 10 different time lags after LC micro-stimulation (from 50-500 ms with 50ms increment steps). In other experiments (n=16) a range of near-threshold stimuli (0, 1/3 T, 2/3 T, T, 4/3 T, 5/3 T, and 2T) were applied to the principal whisker at 5 different time lags from LC micro-stimulation. In each session, deflections were applied in five blocks of 20 pseudo-randomized trials. Across five blocks each test stimulus was presented 100 times with 700-1000 ms random inter-trial interval. For each stimulus amplitude, control trials with no LC micro-stimulation were also intermixed with the main trials. Additionally, there were 100 trials with LC micro-stimulation alone (no-whisker deflection). These trials allowed accurate estimation of the baseline activity at each of the time lags. This procedure was used to control for the fluctuations in the tonic level of NE in the cortex.

### LC micro-stimulation

LC was targeted by a single tungsten microelectrode (<0.7 MΩ, FHC Inc., USA) from 5.6-5.9 mm below the dura. We used a monopolar stimulating electrode to minimize the damage to LC and its projections (Marzo et al., 2014). To confirm the LC location, we applied the following electrophysiological criteria: LC neurons normally have broad extracellular spike waveforms (> 0.6 ms), firing rates of 0.1-6 Hz, and respond to paw pinch with a typical excitation-inhibition pattern (Cedarbaum and Aghajanian, 1978). At the end of the experiment, we verified the LC micro-stimulation site by histology. LC micro-stimulation consisted of a train of 3 bipolar pulses (0.5 ms pulse duration, 100 Hz, 30, 50 or 100 µA). A linear stimulus isolator (A395D, World Precision Instruments, Sarasota, FL, USA) was used to pass current through the electrodes. Due to the technical limitation, we were not able to simultaneously stimulate and record the activity of LC. However, based on previous research currents within 3-300 µA range are expected to activate neurons within a maximal radius of 150 µm (John S. Yeomans, 1990; Snow et al., 1999; Stoney et al., 1968). The precise timing and pattern of micro-stimulation were controlled by MATLAB and fed to the stimulator through the analog output of an NI card (National Instruments, model: BNC 2090, Austin, TX, USA).

### Spike analysis

The sequences of spikes corresponding to trials of the same stimulus were separated and aligned with respect to the stimulus onset to generate raster plots. The occurrence of spiking over time was evaluated by counting the average number of spikes within each time bin. A bin size of 10-20 ms was used to generate prestimulus time histograms (PSTHs). To quantify the effect of LC micro-stimulation on the evoked response, spikes were counted within 1-75 ms and 125-250 ms after deflection (whisker stimulation) for early and late responses, respectively. For spike time analysis, response delay was determined as the first time bin that exceeded background activity by 3 standard deviations (Figure 7).

### Local Field Potential analysis

The raw LFP signals from Barrel Cortex were low pass filtered at 1kHz, and then notch filtered at 60 Hz to remove the city electrical noise. The outlier LFP traces including possible artifacts and saturations were subtracted using the frequency component method (Unakafova and Gail, 2019; Zabeh et al., 2022) which is a conservative cleaning procedure that removes trials with poor signal quality. The power spectrograms of LFP signals were calculated using the wavelet transform, enabling simultaneous localization in the time and frequency domain (van Vugt et al., 2007). The power in each band frequency was subtracted from the trial’s baseline activity during the 200ms pre-stimulation period to cancel out the 1/f color noise (Cohen, 2014).

### Histology

At the end of the experiment, an electrical lesion was made by passing a DC current at 9 V through the LC electrode tip for 10 s. Animals were sacrificed by transcranial perfusion with ∼300 ml saline (0.9%) followed by ∼300 ml phosphate-buffered formalin (10%, pH=7.4). The brain was removed and kept in formalin (for a minimum of one week). Pontine coronal sections (10 µm) were Nissl stained and lesions were detected by light microscopy. LC location was compared with the lesion site using the rat brain atlas (Paxinos and Watson, 2007). For all histological examinations, the micro-stimulation site was anatomically verified to be within LC confirming the reliability of our electrophysiological criteria.

## Results

We electrically stimulated Locus Coeruleus (LC) while recording neuronal activity in the barrel cortex (BC) of urethane anesthetized rats (Figure 1A). LC recording was confirmed based on the characteristic spike waveform, the typical response profile to noxious stimulation, and histology (Figure 1B and C). BC recording was confirmed based on the neuronal response to brief deflections (single-cycle sinusoidal vibration, 12.5 ms duration) applied to the neuron’s principal whisker. At the beginning of each recording session, we applied 10 levels of deflection amplitudes (0-60 μm, 50 repetitions) and fitted a Naka-Rushton function to the average spike count to characterize the neuronal response function and threshold. In the main experiment, we presented a range of deflections centered at the unit’s threshold, T (see Materials and Methods). Figure 1B shows a typical multi-unit activity in three types of trials; control whisker deflection (brown), control LC micro-stimulation (gray), and combined LC stimulation and whisker deflection (blue).

### Modulation of spontaneous activity

LC micro-stimulation exhibited a biphasic effect on BC spontaneous activity: a period of suppression (∼100 ms) followed by a period of excitation (∼200 ms). We applied LC micro-stimulation at three current levels; 30 μA (7 sessions), 50 μA (12 sessions), and 100 μA (21 sessions). At 100 µA current level, micro-stimulation produced a significant reduction of the average spontaneous spiking activity at 1-104 ms followed by an increase in firing rate at 117-325 ms (Figure 2A, top). Across units, maximum inhibition and excitation occurred at 40 ms and 195 ms, respectively. Beyond 325 ms, the spontaneous activity rarely deviated from its pre-stimulation average (Figure 2A, top). The modulation of spontaneous activity was smaller in sessions with 50 µA current level (Figure 2A, middle) and further diminished at 30 µA current level (Figure 2A, bottom). On average, in sessions with 50-µA current, inhibition occurred at 13-108 ms and excitation occurred at 122-309 ms. In 30 μA current sessions, we observed a short period of excitation at 112-143 ms (on average) with no significant inhibition.

**Figure 2.**
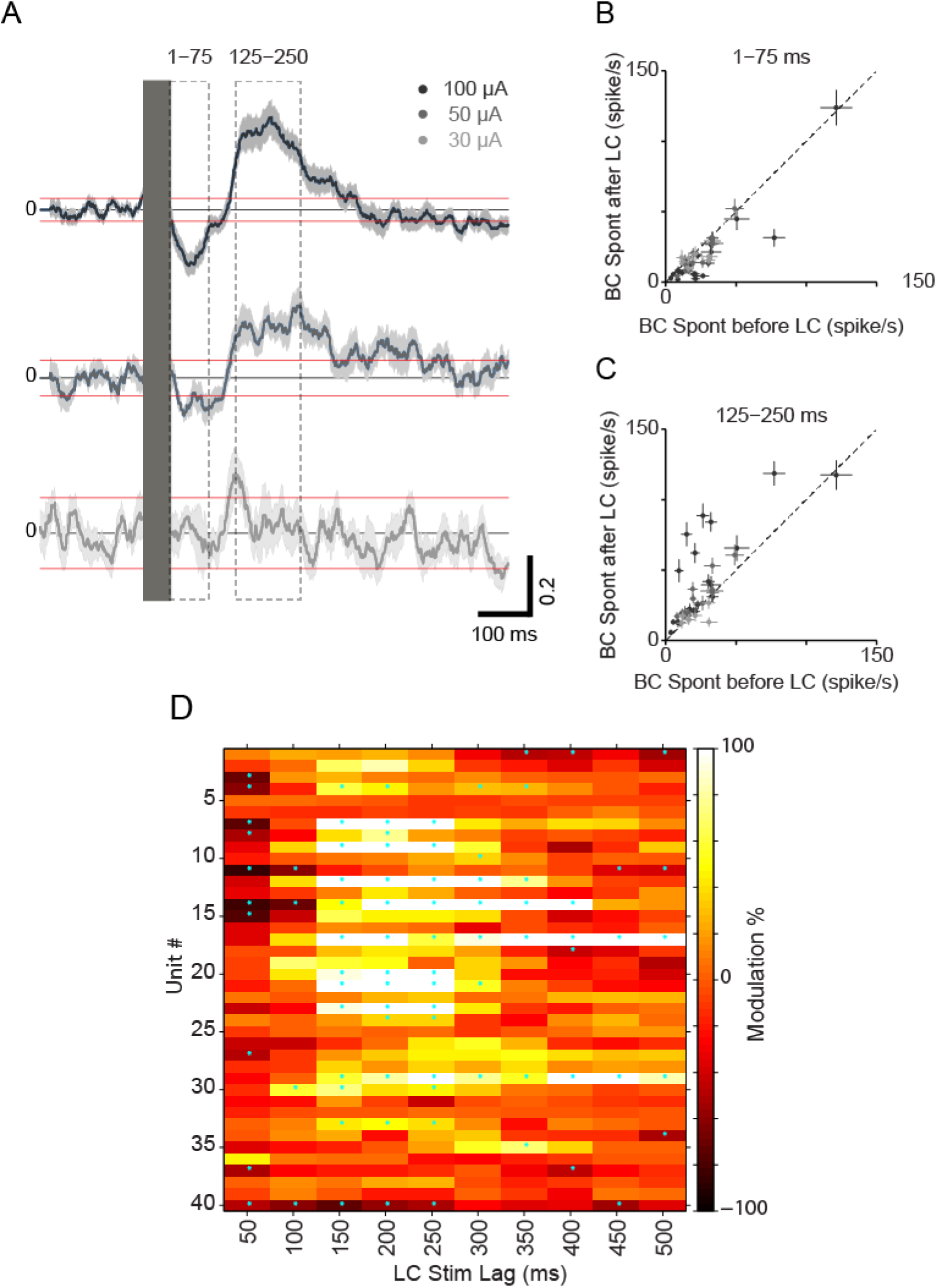
Effect of the LC micro-stimulation on BC spontaneous activity. (A) Average of BC spontaneous activity before and after the LC micro-stimulation (solid box) across multi-units. Each row shows one level of current injection for LC micro-stimulation: 100 µA (top), 50 µA (middle) and 30 µA (bottom). Shaded error bars show standard error of mean (SEM) across multi-units. (B) Every dot represents one multi-unit. Responses show suppression during the early window (1-75 ms post-stimulus onset as outlined in panel A). Error bars show standard error of mean (SEM) across trials for each unit in x and y axes. (C) Same as panel B but for the late excitation phases (125-250 ms, as outlined in panel A). (D) Spontaneous modulation of all units at 10 different time lags.

We further quantified the magnitude of suppression and excitation for individual units (50-µA and 100-µA current levels, n = 33). Based on the profile of modulation at the population level (Figure 2A), we chose two windows of 1-75 and 125-250 ms after LC micro-stimulation to respectively characterize the early and late phases of modulation. The suppression-excitation profile was highly consistent across BC units: 82% of BC units showed early suppression and 91% of BC units showed late excitation (Figure 2B). Overall, the suppression was statistically significant for 70% (23/33) of units and the excitation was statistically significant for 76% (25/33) of units (permutation test, P < 0.05). At 30 μA current level, the suppression and excitation modulation was not as consistent as that observed for the two higher current intensities; although the average firing rate profile did not show a prominent modulation (Figure 2A), 43% (3/7) and 14% (1/7) of units showed significant suppression and excitation (permutation test, P < 0.05) (Figure 2B and C). One multi-unit showed significant suppression at the second window (125-250 ms) in the opposite direction to other units (Figure 2C).

There was no significant correlation between the amount of suppression and the baseline firing rate, indicating that the lack of suppression in some sessions was unlikely to be due to low firing rates (rho = 0.33, P = 0.063, n = 33).

### Modulation of evoked activity

How does LC micro-stimulation modulate the response profile of BC neurons to sensory stimulation? To address this question, we stimulated LC at 10 time lags (50 ms to 500 ms, 50-ms increments) before the application of a single whisker deflection. LC micro-stimulation prominently decreased the early response in the barrel cortex at 50 ms time lag (Figure 3A and B). Overall, 77% (30/39) of units showed a decrease in evoked firing rate (Figure 3B), and this decrease was statistically significant for 60% (100 μA stimulation), 25% (50 μA stimulation), and 14% (30 μA stimulation) of the recorded units (permutation test, P < 0.05). In contrast to early response, the late activity increased in the majority of units, 74% (29/39); 16/20 at 100 µA stimulation, 9/12 at 50 µA stimulation, and 4/7 at 30 µA stimulation (Figure 3C). The enhanced late response was statistically significant for 6/12 units at 50 µA, and 10/20 units at 100 µA (permutation test, P < 0.05).

**Figure 3.**
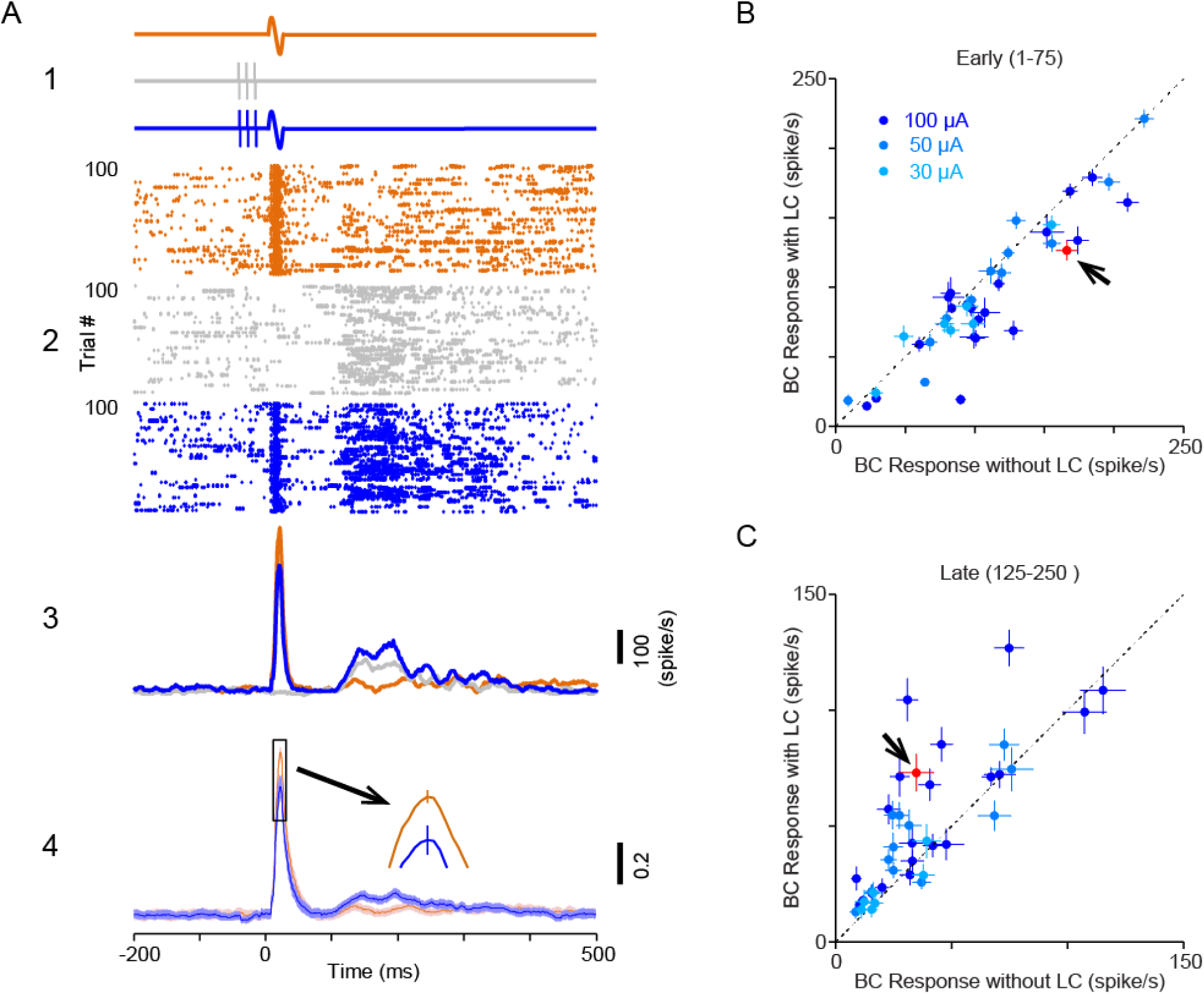
Suppression of BC evoked response by LC micro-stimulation. (A1) LC was micro-stimulated 50-ms before whisker deflection. LC micro-stimulation current was 100 µA. (A2) Raster plot of a typical multi-unit in three types of trials: those with whisker deflections but no LC micro-stimulation (brown); those with LC micro-stimulation but no whisker deflections (gray); and those that contained both LC micro-stimulation and whisker deflections (blue). (A3) PSTHs of the typical unit in A2. (A4) Average of normalized PSTHs across all recorded multi-units. Shaded error bars show standard error of mean (SEM) across multi-units. (B) Scatter plot of the BC early response (1-75) during LC micro-stimulation versus no LC micro-stimulation condition. Each dot-color shows one unit-current level. Error bars show SEM across trials in each axis. A black arrow (red dot) shows a representative unit in panel A. (C) Same as B but for later phase of the evoked response (125-250 ms).

The micro-stimulation effect was highly dependent on the time lag: beyond 50 ms time lag, the early response prominently increased (see for example the 150 ms time lag in Figure 4A and B). Overall, 85% (33/39) of units showed an increase in firing rate (Figure 4B), and this increase was statistically significant for 45% (100 μA stimulation), and 42% (50 μA stimulation) of the recorded units (permutation test, P < 0.05). Figure 4C summarizes the LC evoked modulation at all lags across all units (n = 33) for 100 µA and 50 µA currents.

**Figure 4.**
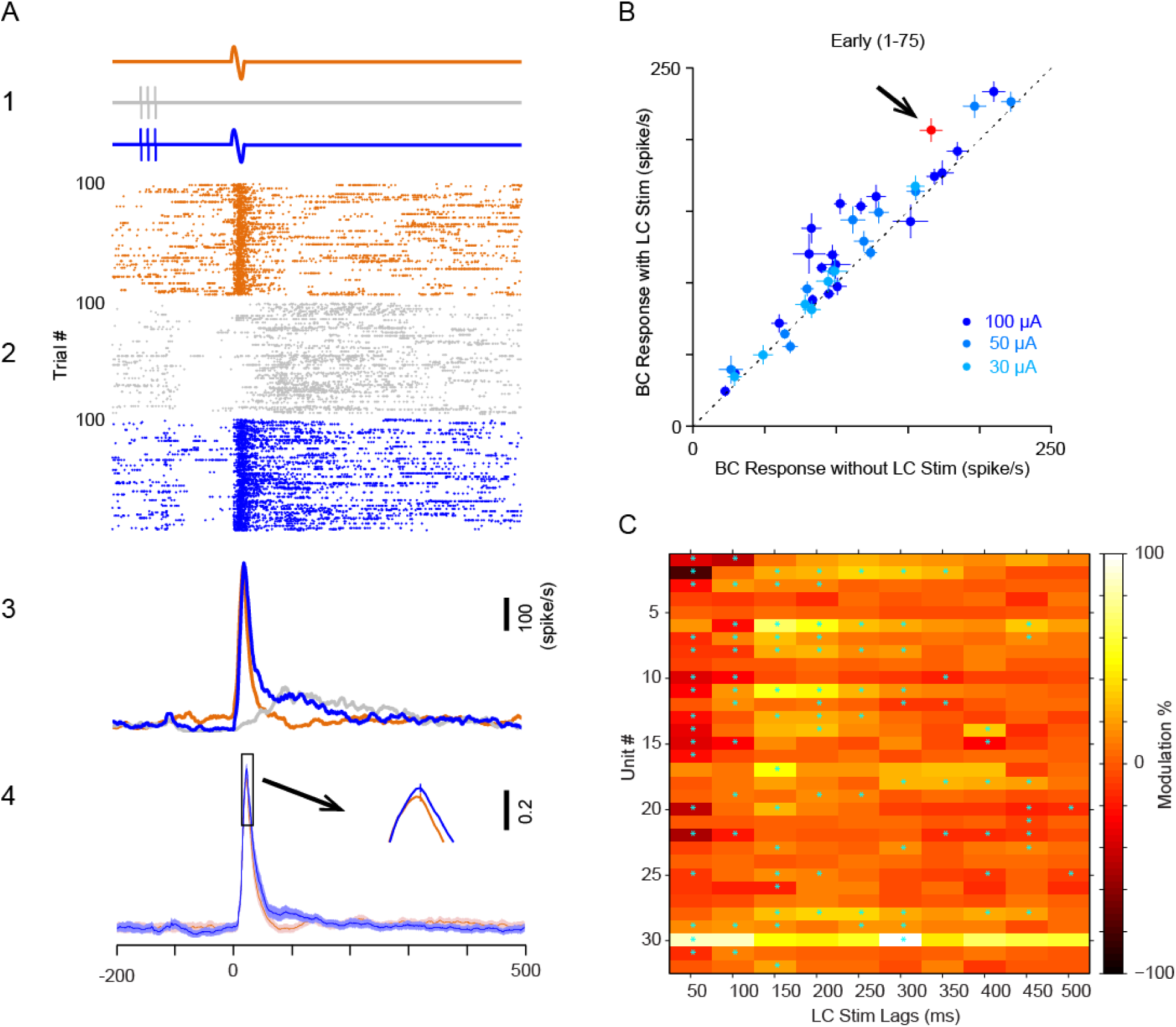
Facilitation of BC evoked responses by LC micro-stimulation. (A1) LC was micro-stimulated 150-ms before whisker deflection. LC micro-stimulation current was 100 µA. (A2) Raster plot of a typical multi-unit in three types of trials: those with whisker deflections but no LC micro-stimulation (brown); those with LC micro-stimulation but no whisker deflections (gray); and those that contained both LC micro-stimulation and whisker deflections (blue). (A3) PSTHs of the typical unit in A2. (A4) Average of normalized PSTHs across all recorded multi-units. Shaded error bars show standard error of mean (SEM) across multi-units. (B) Scatter plot of the BC early response (1-75) during LC micro-stimulation versus no LC micro-stimulation condition. Each dot-color shows one unit-current level. Error bars show SEM across trials in each axis. A black arrow (red dot) shows a representative unit in panel A. (C) Summary of evoked modulation across units in two current levels; 50 and 100 µA (n = 32). Cyan stars show statistically significant modulations, p<0.05, permutation test.

The modulation of the evoked BC response observed in the previous section could simply reflect the modulation of the spontaneous activity as quantified earlier. To examine this possibility, we subtracted for each time lag the peri-stimulus time histogram (PSTH) in the absence of LC micro-stimulation from the one with LC micro-stimulation (Figure 5A). This residual PSTH reflects the changes in firing rate that are produced as a result of LC micro-stimulation (referred to as the Modulation Index in Figure 5C, D). We then overlaid this residual PSTH (black curves) on the spontaneous firing rate (baseline subtracted, gray curves) for each specific time lag (Figure 5A). At 50 ms time lag, the suppression of early-evoked response (∼ 75 ms from deflection onset) was on average higher than the corresponding suppression in spontaneous activity (Figure 5A). The late response (125-250 ms) at 50 ms time lag tended to increase above the changes that were expected based on the spontaneous profile (Figure 5B). At 100 ms time lag, the early suppression was shorter and less pronounced, but the late facilitation was comparable with that of 50 ms time lag. At 150 ms time lag, the early-evoked response was substantially facilitated but the late-evoked response was not significantly modulated compared to the spontaneous activity (Figure 5A). Beyond 150 ms time lag, we observed little modulation of activity except for a slight facilitation in the initial phase of the response (see the 500 ms time lag in Figure 5B). Across all time lags, the activity beyond 700 ms from the deflection onset was similar to the spontaneous activity in the control condition. The modulations at 50 µA current were significantly lower compared to 100 µA (for example for 50-ms lag) but were qualitatively similar. At 30 µA, although some units were affected by LC micro-stimulation, on average the modulation pattern was absent.

**Figure 5.**
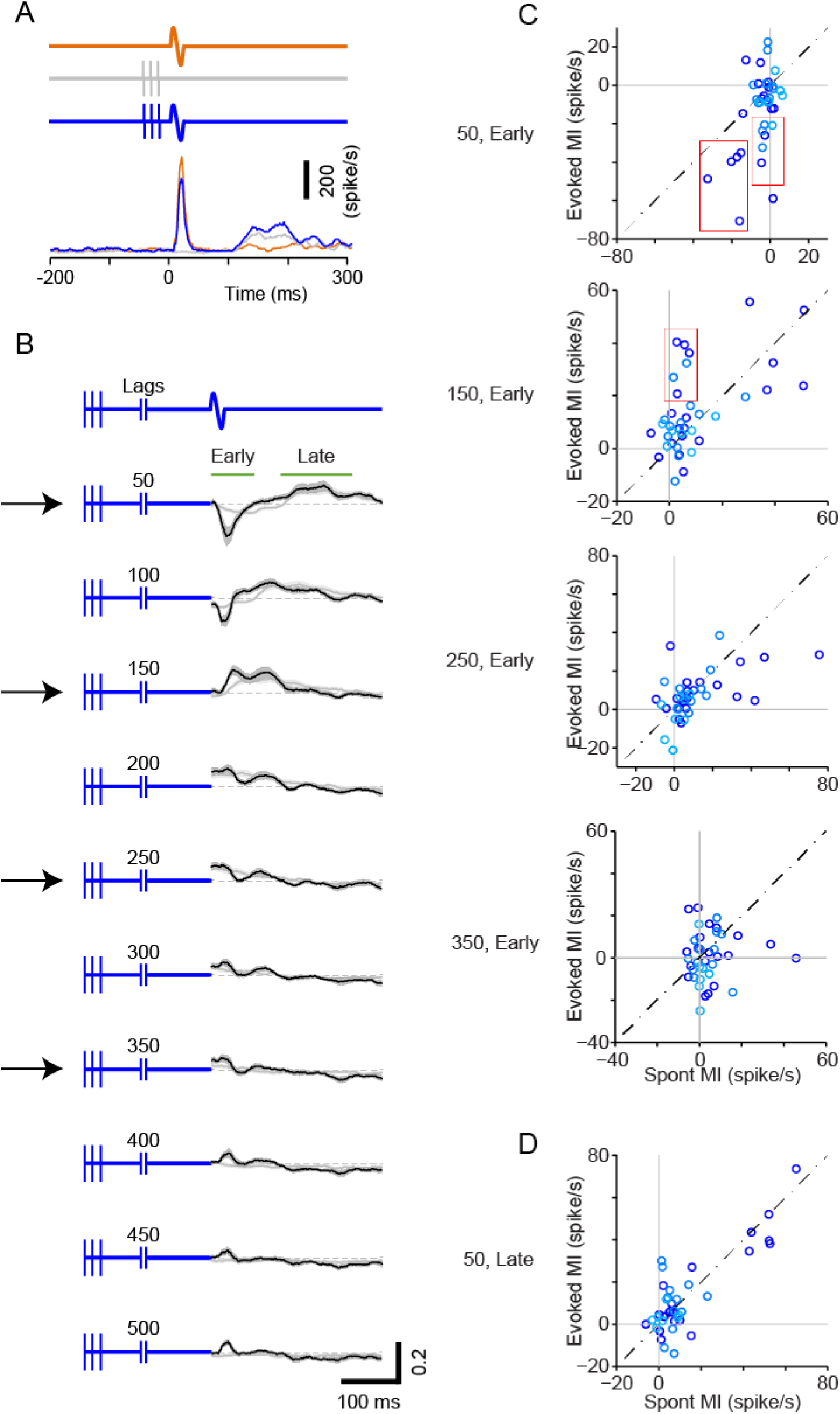
LC micro-stimulation modulation of spontaneous and evoked activity. (A) PSTH a representative unit. (B) Temporal profile of LC modulation of BC activity; spontaneous and evoked. Top row shows the time course of LC and whisker stimulation across various trial types. LC micro-stimulation was applied at different time lags from 50 to 500 ms with a 50-ms step. Gray and black traces show an average of normalized spontaneous and evoked modulation, respectively. Each unit was normalized to peak firing rate in the first 50 ms after stimulus onset. Spontaneous modulation is the change in activity after LC micro-stimulation in the absence of any whisker. Evoked modulation is the difference of deflection-evoked PSTHs between LC micro-stimulation and no-LC micro-stimulation trials. The vertical blue lines show the timing of LC micro-stimulation. Dotted lines show 0 point of modulation. Each trace shows the average of modulation across all recorded units for 100 µA current level. Three arrows show three lags in panel C. (C) Scatter plots show evoked modulation versus spontaneous modulation for the early response (1:75) in four lags that exhibited the highest modulation. Each dot shows one unit. Dark blue represents 100 µA stimulation, light blue represents 50 µA stimulation. Dashed lines show unity lines. (D) Same as C but for the late response (125-250) at 50 ms lag.

Finally, we asked whether the modulation of spontaneous activity could explain the modulation of evoked response across recording sessions. Figure 5B plots the spontaneous modulation index against the early-evoked modulation index (1-75 ms from the deflection onset). At 50 ms time lag across the 3 current levels, 77% of the sessions showed higher evoked modulation than spontaneous (n=30/39). Notably, the sessions with little modulation of spontaneous activity had comparable (permutation test, P = 0.27) baseline firing rate with those which had stronger spontaneous modulation (red boxes in figure 5C), indicating that the difference between evoked and spontaneous modulation was not due to a floor effect when baseline activity might be too low to exhibit suppression of firing rate. At 150 ms lag, the evoked modulation was higher than spontaneous modulation in 64% of the sessions (n = 25/39). For 5 out of 6 units outlined with a red box in Figure 5C, the spontaneous activity was not significantly modulated (permutation test), while the evoked response was substantially enhanced (Figure 5C, 150 early). In contrast, in 250 ms lag, 64% of the units showed higher spontaneous modulation indicating a relative suppression of the evoked response. Unlike the early response, for the majority of units, the late activity did not show a significant modulation above the spontaneous profile: at 50-ms time lag, for the 125-250 ms post deflection onset, modulation of evoked activity paralleled that of spontaneous activity (note that most units are dispersed along the diagonal line in Figure 5C). This is consistent with a linear summation of the LC-evoked spikes and deflection-evoked spikes and hence no net modulation of the evoked response.

### Modulation of response latency

The average profile of modulation in Figure 5 suggests that LC micro-stimulation can affect the temporal structure of the response. For example, in 50 and 100 ms time lags, we observed a suppression followed by facilitation in the modulation profile (Figure 5B). We, therefore, asked if LC stimulation affected the neuronal response latencies to sensory stimulation.

Figure 6A quantifies the modulation of response latency across all time lags (see Materials and methods). Although the average response latency appeared to increase at 100 ms lag, this change was not statistically significant (P = 0.055; control condition, 12.8 ± 0.6 and LC micro-stimulation, 13.9 ± 0.6; Figure 6B). However, the coupling of deflection with LC micro-stimulation at 250 ms time lag significantly decreased the response latency from 12.8 ± 0.6 to 11.8 ± 0.6 (P < 0.001, permutation test, Figure 6C).

**Figure 6.**
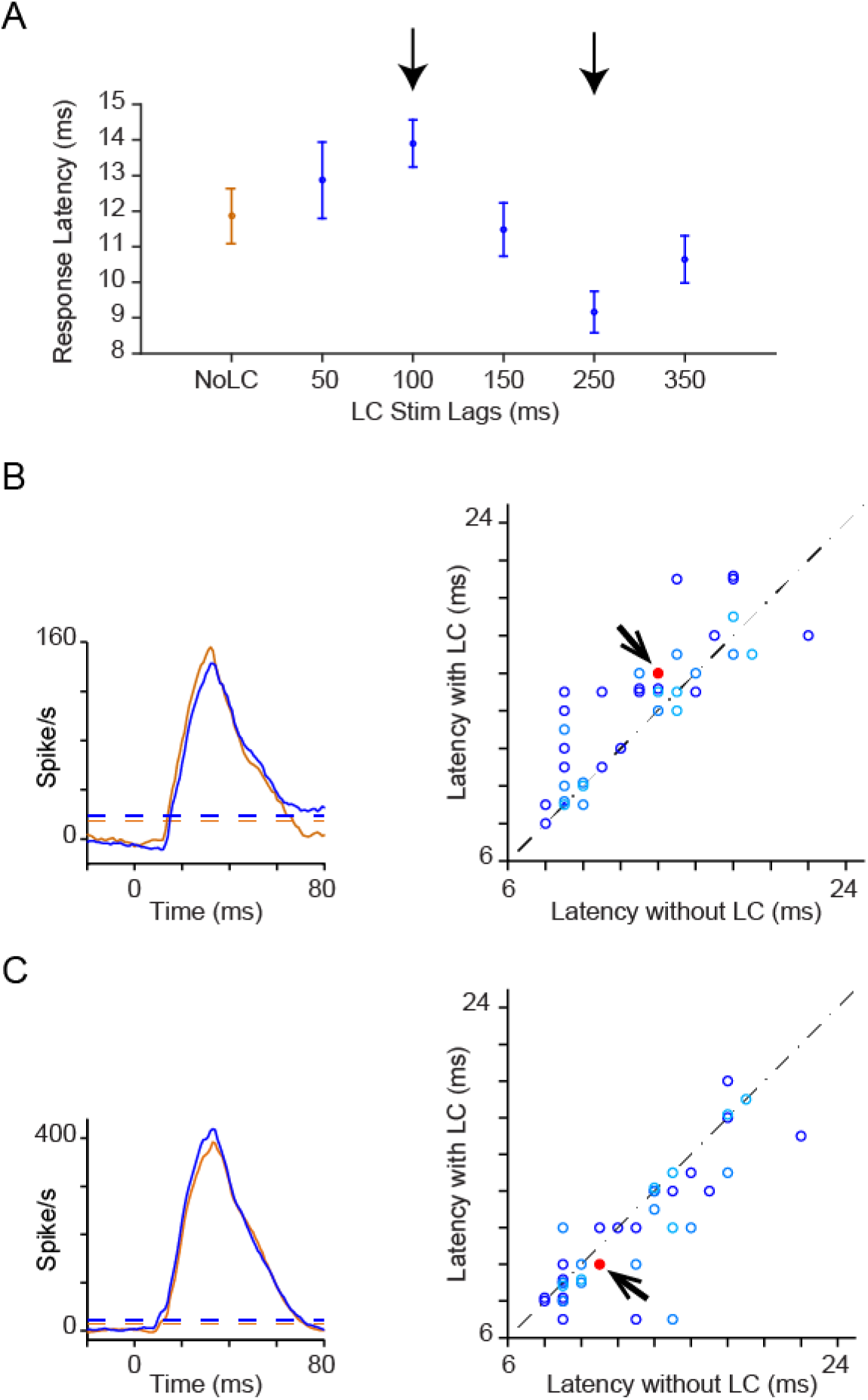
Effect of LC micro-stimulation on response latency. (A) Average response latency across all units and current levels. At 100 ms lag, the latency significantly increased and at 250 ms lag, the latency significantly decreased (p<0.05, permutation test). (B) Left; PSTHs of a representative multi-unit to whisker deflection of threshold (T) amplitude (bin size, 20 ms; sliding at 1-ms steps). Micro-stimulation was at 100 ms lag. Right; response latency for all units which showed significant response in two conditions (n = 36). Red dots in B and C show a representative unit in the left panel. (C) Left; PSTHs of another representative multi-unit. LC micro-stimulation was at 250 ms lag. Right; response latency for all units which showed significant response in two conditions (n = 36).

### Modulation of the stimulus-response function

How does LC micro-stimulation affect the stimulus-response function and the sensitivity of neurons in detecting sensory stimuli? To address this question, in a subset of experiments we applied a range of near-threshold stimuli; 1/3 T, 2/3 T, T, 4/3 T, 5/3 T, and 2T and obtained a full stimulus-response function in the presence and absence of LC activation (Figure 7). LC micro-stimulation at 50 ms lag decreased the neuronal response at all stimulus intensities (Figure 7A, top). On the other hand, at 150 ms lag, LC micro-stimulation increased the response at all stimulus intensities (Figure 7A, middle). This enhancement of response diminished at 250 ms lag (Figure 7A, bottom). The overall upward shift in the response function resulted in a reduced response range (Figure 7B).

**Figure 7.**
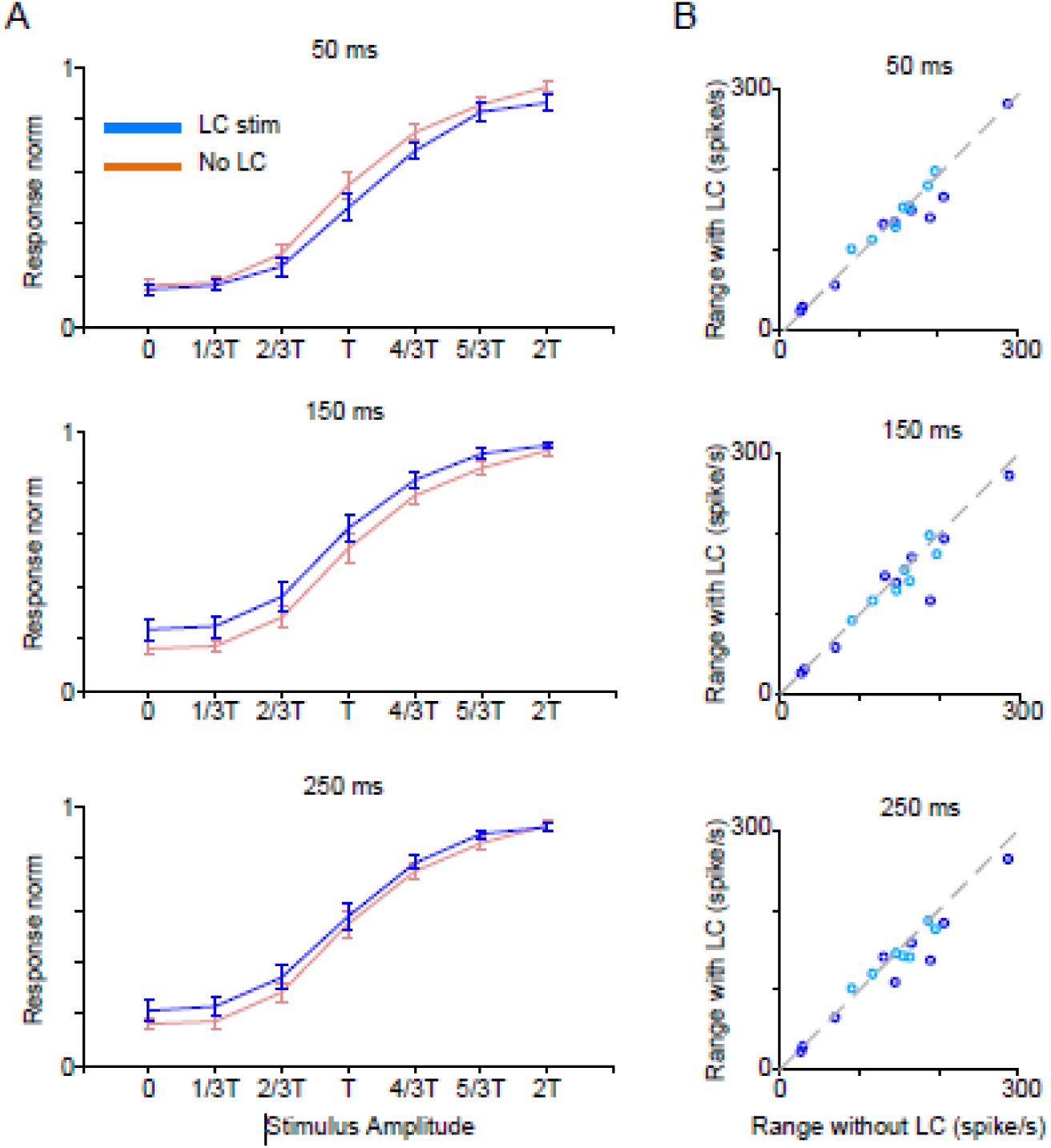
Effect of LC micro-stimulation on stimulus-response function. (A) The average tuning function across all recorded units at two levels of current (30 and 100 µA) in response to the full range of stimulus amplitudes (n = 16). (B) Response range was calculated as the difference between the highest and lowest response for each unit. Blue and light blue circles show data for 100 µA and 30 µA respectively. Each panel shows data for three LC micro-stimulation lags; 50, 150, and 250 ms. Error bars are SEM across units. Spike count was calculated over a 75-ms window post-stimulus onset.

### Modulation of cortical state

Finally, we quantified how LC micro-stimulation modulated the overall state of the cortex. The change in cortical state can be quantified using low-frequency oscillations of EEG and LFP signals (Fazlali et al., 2016; Harris and Thiele, 2011). We, therefore, analyzed LFP recorded from the barrel cortex to quantify the effect of LC stimulation on the cortical state. The average power spectrogram of LFP indicates that LC stimulation suppressed the low-frequency power (Figure 8A). Across sessions, the theta oscillation power (4–8 Hz) was suppressed while there were no significant power changes for higher frequency oscillations such as Gamma (30-50 Hz), (Figure 8B; Wilcoxon rank sum test P=0.1 for 100 mA, P = 0.9 for 50mA). The effect of stimulation on cortical oscillation initiated beyond 200 ms after stimulation onset and continued beyond 1 second. The 30 mA LC stimulation did not initiate significant modulations in the BC oscillations.

**Figure 8.**
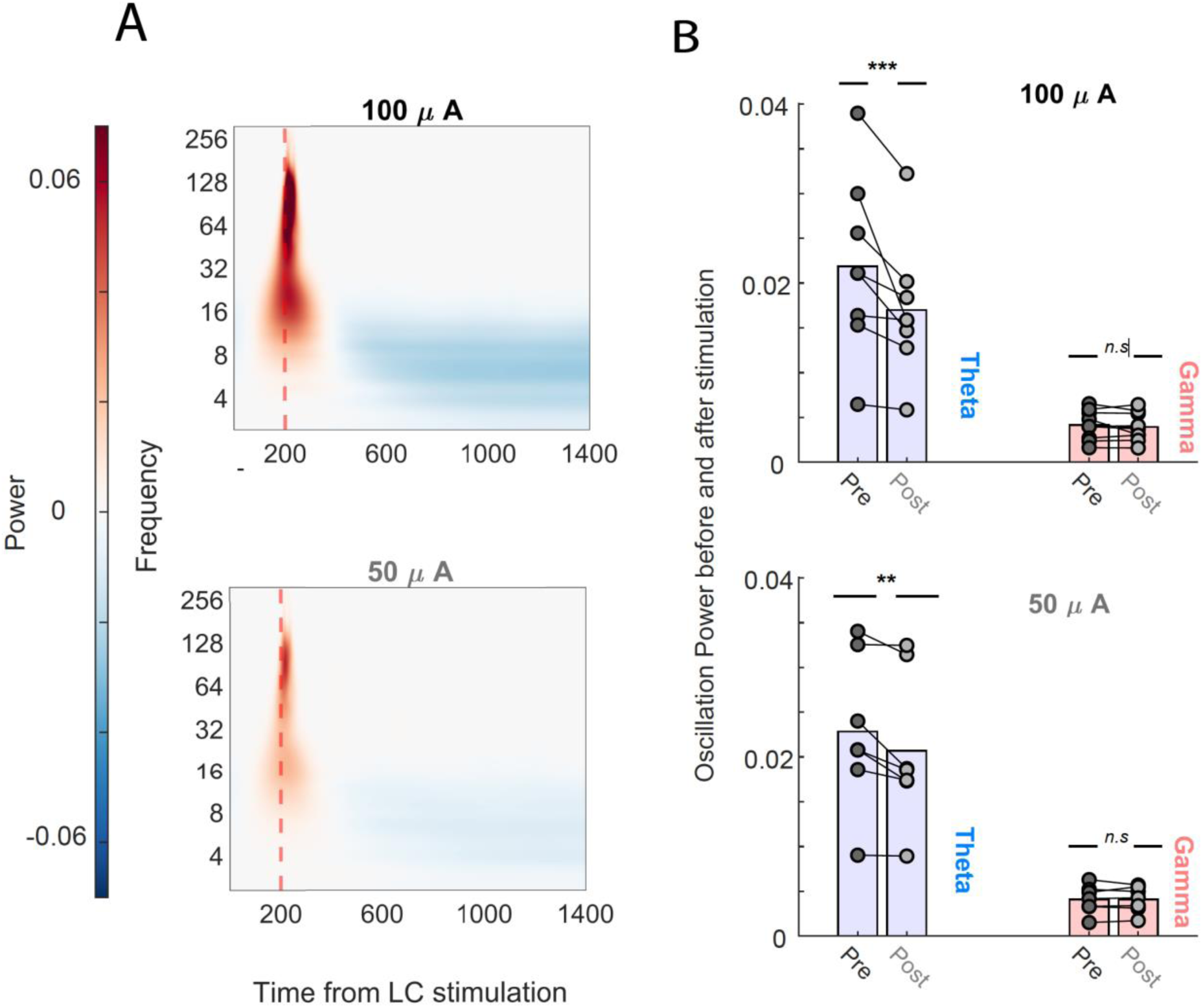
Effect of LC stimulation on the brain state: (A) Average cortical power spectrogram at two levels of LC stimulation current (50 and 100 µA) across 100 trials of an example session. Low frequency (alpha-theta) oscillations were suppressed after 200 ms of stimulation onset. (B) Across session comparison of pre and post-stimulation power of Theta and Gamma oscillations at two levels of LC stimulation current (50 and 100 µA). Each line represents one session and each dot represents trial average oscillation power during 0 -150ms pre-stimulation, and 1000-1150ms post-stimulation. ***P<0.01, **P<0.05, Wilcoxon rank-sum test.

## Discussion

Here, we studied the temporal profile of LC modulation of the sensory processing by coupling spontaneous and sensory input-driven activity in the rat barrel cortex with phasic LC activation across a range of time lags (50-500 ms). LC micro-stimulation exhibited a biphasic effect on BC spontaneous activity: a period of suppression followed by a period of excitation consistent with earlier experiments (Phillis and Kostopoulos, 1977). The effect of LC micro-stimulation on the evoked response to the near-threshold stimulus was highly time-dependent: at 50-ms lag, the early-evoked response decreased in most of the units. This contrasted with 150-ms lag, where a majority of units exhibited response facilitation. At 150 to 350-ms time lags, LC micro-stimulation caused a combined facilitation followed by suppression. At higher time lags (≥400), the effect was mainly facilitation. Response latency was decreased at 250 ms following the LC micro-stimulation.

### LC micro-stimulation and sensory processing

The effect of LC micro-stimulation on sensory responses is often quantified in time lags between 250 to 350 ms (Bouret and Sara, 2002; Edeline et al., 2011; Lecas, 2004). LC responds to sensory stimulation with a short latency in awake animals (Aston-Jones and Bloom, 1981b; Foote et al., 1980). A previous study compared the modulation of sensory response in the somatosensory cortex at time intervals of 100-500 ms and found mainly potentiation with a peak at 200 ms (Waterhouse et al., 1998). However, micro-stimulation parameters were adjusted to be sub-threshold for modulation of spontaneous activity (Lecas, 2004; Lecas et al., 2001; Waterhouse et al., 1998). In our study, different current levels were used to compare the relationship between the modulation of spontaneous and evoked activities. Our data suggest that although the modulation of sensory response by LC micro-stimulation lasts for a long period (> 500ms), the peak of effects can emerge within 100 ms as a strong suppression of both spontaneous and evoked activity, and this is followed by facilitation at 150 ms time lag. We found various degrees of correlation between spontaneous and evoked activity; in some sessions, evoked activity was modulated independently of modulation in spontaneous activity (Figure 5B). We observed less modulation in evoked activity when the current was adjusted to have a minimal effect on spontaneous activity (30 µA, Figure 5B).

### Effectiveness and specificity of LC activation

The level of brain tissue activation by an electrical current depends on multiple parameters such as current pulse amplitude, polarity, duration, and frequency (McIntyre and Grill, 2000; Tehovnik et al., 2006). Since myelinated axon fibers have higher excitability (Nowak and Bullier, 1998; Tehovnik et al., 2006), the spread of current in pons may directly activate the surrounding fibers. This can affect other cortical areas with a chain of trans-synaptic activations/inhibitions resulting in the observed effects. Consistent with this possibility, we observed bilateral facial movements with 100 µA current in several recording experiments (data not included). With lower currents (50 and 30 µA), no facial movement was observed. A study reported that this level of stimulation (30 and 50 µA, 0.4 ms duration) can produce bilateral modulation of LC activity (Marzo et al., 2014). Likewise, in our study, the direct or indirect activation of other structures, including pontine reticular formation and basal forebrain (Dringenberg and Vanderwolf, 1997) cannot be ruled out. Future optogenetic experiments can directly address this concern.

We presented a range of whisker stimuli to quantify the effect of LC micro-stimulation on cortical stimulus-response functions. At least one study has shown that phasic LC stimulation, 300 ms before whisker deflection, causes a leftward shift in stimulus-response function and greatly facilitates response to weaker stimuli in the thalamus (Berridge and Waterhouse, 2003; Vazey et al., 2018; Yang et al., 2021). In our data, a suppressive effect at 50-ms lag and a facilitation at 150-250 ms was consistently observed for near-threshold stimuli. On the other hand, the response to the highest stimulus intensity (2T) remained unchanged with LC micro-stimulation (Figure 7A). We found that strong LC micro-stimulation is likely to affect near-threshold stimuli, decrease the range of responsiveness (Figure 7B) and disrupt the encoding (McBurney-Lin et al., 2019; McCormick et al., 2020; McGinley et al., 2015).

### LC activity and brain state

The role of LC-NE system in the modulation of the cortical state is implicated in several studies (Durán et al., 2021; Hayat et al., 2020; Marzo et al., 2014; Noei et al., 2020; Zagha and McCormick, 2014). There is a strong correlation between the tonic activity of the LC and two extreme cortical states; the low LC firing rate is associated with the synchronized state and the high LC firing rate with the desynchronized state (Aston-Jones and Bloom, 1981a; Fazlali et al., 2016; Safaai et al., 2015). The suppression of low-frequency oscillation is associated with the brain state shift toward higher levels of arousal (Fazlali et al., 2016; Harris and Thiele, 2011; Lee et al., 2020; Weiss et al., 2023).

We showed that the effect of LC stimulation on cortical oscillation was exclusive to theta oscillation (low frequency) and it did not affect the higher frequencies such as the gamma oscillation (Figure 8B). Earlier electrophysiology studies across species have shown that the Theta oscillation is dominant during REM sleep, whereas Gamma is associated with coordinated interaction of local network during active processing and behavior (Buzśaki and Wang, 2012; Jacobs, 2014)

Our results indicate that the LC modulates both the spiking activity of sensory-driven activity of the neurons and cortical oscillations, each with a different temporal profile. LC modulations of state initiated after 200 ms and continued for more than 1 second. This was at a slower time course than that observed in LC modulations of spiking (i.e. 50-350 ms after LC stimulation). This different temporal effect of LC stimulation can be explained by the activation of a distinct population of LC neurons (Breton-Provencher and Sur, 2019; Poe et al., 2020; Totah et al., 2018). The later effect of LC stimulation on the cortical state can be explained by the indirect effects of LC activation and the interplay between various neuromodulatory systems (Lee and Dan, 2012; Ranjbar-Slamloo and Fazlali, 2020; Sara, 2009) and thalamus (Hirata and Castro-Alamancos, 2010; Poulet et al., 2012; Rodenkirch et al., 2019) for changing the brain state (McGinley et al., 2015; Sabri and Arabzadeh, 2018; Weiss et al., 2023; Zagha and McCormick, 2014).

### LC topography, projections, and their effect on BC spontaneous activity

The LC micro-stimulation with 100 and 50 µA currents revealed a biphasic effect on spontaneous activity; an early inhibition followed by an excitation (Figure 2A). LC neurons projecting to cortical and subcortical areas are not homogenously distributed across this nucleus (Bari et al., 2020; Breton-Provencher and Sur, 2019; Chandler et al., 2019; Poe et al., 2020; Sara and Bouret, 2012; Schwarz and Luo, 2015). Cortical projecting neurons are located at the body, while thalamic projecting neurons are located in the posterior tail of LC (Loughlin et al., 1986; Schwarz and Luo, 2015). Through dense projections to the thalamic reticular nucleus (TRN), LC can gate information flow in the whisker pathway (Rodenkirch et al., 2019). The high variability that we observed across sessions might be due to session-to-session variability in targeting different parts of LC (Breton-Provencher and Sur, 2019; Chandler et al., 2019; Totah et al., 2018, 2019). Moreover, the biphasic nature of the cortical modulation caused by larger currents might be due to the activation of separate LC-NE projecting pathways (Logothetis et al., 2010).

### LC activity and behavior

The central nervous system’s ability to efficiently extract relevant information from the sensory environment is essential for survival. The whisker system is one of the main channels through which rodents collect information from their environment (Diamond and Arabzadeh, 2013). Behavioral studies have revealed a tight correlation between neuronal activity in the barrel cortex and whisker-mediated behavior (O’Connor et al., 2013, 2010; von Heimendahl et al., 2007). These behavioral studies typically use well-trained animals with reliable levels of performance. The degree of synchronization in cortical cells is shown to depend on the level of training: naïve animals elicit a more synchronized cortical state compared to trained animals (Sachidhanandam et al., 2013). The LC-NE modulation of the cortical state may play a role in improved performance observed with training. Consistent with this idea, the activity of the LC-NE system and the tightly coupled pupil dynamics are shown to predict the behavioral performance of rodents (Lee et al., 2020; Schriver et al., 2020) and primates (Aston-Jones et al., 1999; Aston-Jones and Cohen, 2005).

### Concluding remarks

The majority of studies on LC-NE system and sensory properties suggest a role for this system in the processing of salient stimuli and perceptual acuity (Aston-Jones and Cohen, 2005; Sara, 2009; Waterhouse and Navarra, 2019). Our data suggest that the time lag of coupling between LC activation and sensory input is a crucial parameter for the interpretation of its neuromodulatory effects. In addition to studies that show the diverse population of LC neurons (Loughlin et al., 1986; Schwarz and Luo, 2015), there is growing evidence pointing to the subpopulation of LC neurons with different electrophysiology characteristics (Poe et al., 2020; Totah et al., 2018) implicated in attention (Bari et al., 2020), arousal (Breton-Provencher and Sur, 2019) and brain state (Noei et al., 2020). Our results were consistent with these findings and indicate that the effect of LC activation on information processing and brain state works at multiple time courses.

## Abbreviations

(LC): Locus Coeruleus
(BC): Barrel Cortex
(NE): Norepinephrine
(LFP): Local Field Potential

## Author Contributions

ZF, YR-S, and EA designed research; ZF and YR-S performed experiments; ZF analyzed the data; ZF prepared figures 1-7; ZF and EZ analyzed brain oscillation section, figure 8; ZF, YR-S, EZ and EA interpreted results of experiments; ZF drafted manuscript; EA supervised study; ZF, YR-S, EZ and EA read and revised the manuscript.

## Data Availability Statement

The data that support the findings of this study are available on request from the corresponding author.

## Acknowledgements

We are grateful to Hossein Esteky, Behrad Noudoost and Saeed Semnanian for providing technical support and helpful discussions.

## References

Aston-Jones G, Bloom FE. 1981a. Activity of norepinephrine-containing locus coeruleus neurons in behaving rats anticipates fluctuations in the sleep-waking cycle. J Neurosci 1:876–886.

Aston-Jones G, Bloom FE. 1981b. Norepinephrine-containing locus coeruleus neurons in behaving rats exhibit pronounced responses to non-noxious environmental stimuli. J Neurosci 1:887–900.

Aston-Jones G, Cohen JD. 2005. An integrative theory of locus coeruleus-norepinephrine function: adaptive gain and optimal performance. Annu Rev Neurosci 28:403–50. doi:10.1146/annurev.neuro.28.061604.135709

Aston-Jones G, Rajkowski J, Cohen J. 1999. Role of Locus Coeruleus in Attention and Behavioral Flexibility. Biol Psychiatry 46:1309–1320.

Bari A, Xu S, Pignatelli M, Takeuchi D, Feng J, Li Y, Tonegawa S. 2020. Differential attentional control mechanisms by two distinct noradrenergic coeruleo-frontal cortical pathways. Proc Natl Acad Sci U S A 117:29080–29089. doi:10.1073/pnas.2015635117

Berridge CW, Waterhouse BD. 2003. The locus coeruleus-noradrenergic system: modulation of behavioral state and state-dependent cognitive processes. Brain Res Rev 42:33–84.

Bouret S, Sara SJ. 2004. Reward expectation, orientation of attention and locus coeruleus-medial frontal cortex interplay during learning. Eur J Neurosci 20:791–802. doi:10.1111/j.1460-9568.2004.03526.x

Bouret S, Sara SJ. 2002. Locus coeruleus activation modulates firing rate and temporal organization of odour-induced single-cell responses in rat piriform cortex. Eur J Neurosci 16:2371–2382. doi:10.1046/j.1460-9568.2002.02413.x

Breton-Provencher V, Sur M. 2019. Active control of arousal by a locus coeruleus GABAergic circuit. Nat Neurosci 22:218–228. doi:10.1038/s41593-018-0305-z

Buzśaki G, Wang XJ. 2012. Mechanisms of gamma oscillations. Annu Rev Neurosci 35:203– 225. doi:10.1146/annurev-neuro-062111-150444

Carter ME, Yizhar O, Chikahisa S, Nguyen H, Adamantidis A, Nishino S, Deisseroth K, de Lecea L. 2010. Tuning arousal with optogenetic modulation of locus coeruleus neurons. Nat Neurosci 13:1526–33. doi:10.1038/nn.2682

Cedarbaum JM, Aghajanian GK. 1978. Activation of locus coeruleus neurons by peripheral stimuli: modulation by a collateral inhibitory mechanism. Life Sci 23:1383–1392.

Chandler DJ, Jensen P, McCall JG, Pickering AE, Schwarz LA, Totah NK. 2019. Redefining Noradrenergic Neuromodulation of Behavior: Impacts of a Modular Locus Coeruleus Architecture. J Neurosci 39:8239–8249. doi:10.1523/JNEUROSCI.1164-19.2019

Cohen MX. 2014. Analyzing Neural Time Series Data. Anal Neural Time Ser Data. doi:10.7551/mitpress/9609.001.0001

Devilbiss DM, Page ME, Waterhouse BD. 2006. Locus ceruleus regulates sensory encoding by neurons and networks in waking animals. J Neurosci 26:9860–72. doi:10.1523/JNEUROSCI.1776-06.2006

Devilbiss DM, Waterhouse BD. 2011. Phasic and tonic patterns of locus coeruleus output differentially modulate sensory network function in the awake rat. J Neurophysiol 105:69–87. doi:10.1152/jn.00445.2010

Devilbiss DM, Waterhouse BD. 2004. The effects of tonic locus ceruleus output on sensory-evoked responses of ventral posterior medial thalamic and barrel field cortical neurons in the awake rat. J Neurosci 24:10773–85. doi:10.1523/JNEUROSCI.1573-04.2004

Diamond ME, Arabzadeh E. 2013. Whisker sensory system – From receptor to decision. Prog Neurobiol 103:28–40. doi:http://dx.doi.org/10.1016/j.pneurobio.2012.05.013

Dringenberg HC, Vanderwolf CH. 1997. Neocortical activation: modulation by multiple pathways acting on central cholinergic and serotonergic systems. Exp Brain Res 116:160–174. doi:10.1007/PL00005736

Durán E, Yang M, Neves R, Logothetis NK, Eschenko O. 2021. Modulation of Prefrontal Cortex Slow Oscillations by Phasic Activation of the Locus Coeruleus. Neuroscience 453:268–279. doi:10.1016/j.neuroscience.2020.11.028

Edeline J-M. 2012. Beyond traditional approaches to understanding the functional role of neuromodulators in sensory cortices. Front Behav Neurosci 6:45. doi:10.3389/fnbeh.2012.00045

Edeline J-M, Manunta Y, Hennevin E. 2011. Induction of selective plasticity in the frequency tuning of auditory cortex and auditory thalamus neurons by locus coeruleus stimulation. Hear Res 274:75–84. doi:10.1016/j.heares.2010.08.005

Eldar E, Cohen JD, Niv Y. 2013. The effects of neural gain on attention and learning. Nat Neurosci 16:1146–1153. doi:10.1038/nn.3428

Fazlali Z, Ranjbar-Slamloo Y. 2021. Locus Coeruleus Stimulation Affects response Adaptation in the Somatosensory Cortex of Whisker-to-barrel Touch System. bioRxiv 2021.05.19.444702. doi:10.1101/2021.05.19.444702

Fazlali Z, Ranjbar-Slamloo Y, Adibi M, Arabzadeh E. 2016. Correlation between cortical state and locus coeruleus activity: Implications for sensory coding in rat barrel cortex. Front Neural Circuits 10. doi:10.3389/fncir.2016.00014

Foote SL, Aston-Jones G, Bloom FE. 1980. Impulse activity of locus coeruleus neurons in awake rats and monkeys is a function of sensory stimulation and arousal. Proc Natl Acad Sci U S A 77:3033–3037.

Foote SL, Freedman R, Oliver AP. 1975. Effects of putative neurotransmitters on neuronal activity in monkey auditory cortex. Brain Res 86:229–242. doi:http://dx.doi.org/10.1016/0006-8993(75)90699-X

Gottlieb J, Cohanpour M, Li Y, Singletary N, Zabeh E. 2020. Curiosity, information demand and attentional priority. Curr Opin Behav Sci 35:83–91. doi:10.1016/j.cobeha.2020.07.016

Harris KD, Thiele A. 2011. Cortical state and attention. Nat Rev Neurosci 12:509–23. doi:10.1038/nrn3084

Hayat H, Regev N, Matosevich N, Sales A, Paredes-Rodriguez E, Krom AJ, Bergman L, Li Y, Lavigne M, Kremer EJ, Yizhar O, Pickering AE, Nir Y. 2020. Locus coeruleus norepinephrine activity mediates sensory-evoked awakenings from sleep. Sci Adv 6. doi:10.1126/sciadv.aaz4232

Hirata A, Aguilar J, Castro-Alamancos M a. 2006. Noradrenergic activation amplifies bottom-up and top-down signal-to-noise ratios in sensory thalamus. J Neurosci 26:4426–36. doi:10.1523/JNEUROSCI.5298-05.2006

Hirata A, Castro-Alamancos MA. 2010. Neocortex network activation and deactivation states controlled by the Thalamus. J Neurophysiol 103:1147–1157. doi:10.1152/jn.00955.2009

Jacobs J. 2014. Hippocampal theta oscillations are slower in humans than in rodents: Implications for models of spatial navigation and memory. Philos Trans R Soc B Biol Sci 369. doi:10.1098/rstb.2013.0304

Jiang M, Griff ER, Ennis M, Zimmer LA, Shipley MT. 1996. Activation of Locus Coeruleus Enhances the Responses of Olfactory Bulb Mitral Cells to Weak Olfactory Nerve Input 16:6319–6329.

John S. Yeomans. 1990. Principles of Brain Stimulation, Oxford University Press. doi:10.1097/00004691-199201000-00038

Lecas J. 2004. Locus coeruleus activation shortens synaptic drive while decreasing spike latency and jitter in sensorimotor cortex . Implications for neuronal integration. Eur J Neurosci 19:2519–2530. doi:10.1111/j.1460-9568.2004.03341.x

Lecas J, Neuromodulation D, Neurosciences I, Pierre U, Bât B. 2001. Noradrenergic modulation of tactile responses in rat cortex . Current source-density and unit analyses 324:33–44.

Lee CCY, Kheradpezhouh E, Diamond ME, Arabzadeh E. 2020. State-Dependent Changes in Perception and Coding in the Mouse Somatosensory Cortex. Cell Rep 32:108197. doi:10.1016/j.celrep.2020.108197

Lee S-H, Dan Y. 2012. Neuromodulation of brain states. Neuron 76:209–22. doi:10.1016/j.neuron.2012.09.012

Logothetis NK, Augath M, Murayama Y, Rauch A, Sultan F, Goense J, Oeltermann A, Merkle H. 2010. The effects of electrical microstimulation on cortical signal propagation. Nat Neurosci 13:1283–1291. doi:10.1038/nn.2631

Loughlin SE, Foote SL, Bloom FE. 1986. Efferent projections of nucleus locus coeruleus: Topographic organization of cells of origin demonstrated by three-dimensional reconstruction. Neuroscience 18:291–306. doi:10.1016/0306-4522(86)90155-7

Manunta Y, Edeline J-M. 2004. Noradrenergic induction of selective plasticity in the frequency tuning of auditory cortex neurons. J Neurophysiol 92:1445–63. doi:10.1152/jn.00079.2004

Marzo A, Totah NK, Neves RM, Logothetis NK, Eschenko O. 2014. Unilateral electrical stimulation of rat locus coeruleus elicits bilateral response of norepinephrine neurons and sustained activation of medial prefrontal cortex. J Neurophysiol 111:2570–88. doi:10.1152/jn.00920.2013

McBurney-Lin J, Lu J, Zuo Y, Yang H. 2019. Locus coeruleus-norepinephrine modulation of sensory processing and perception: A focused review. Neurosci Biobehav Rev 105:190– 199. doi:10.1016/j.neubiorev.2019.06.009

McCormick DA, Nestvogel DB, He BJ. 2020. Neuromodulation of Brain State and Behavior. Annu Rev Neurosci 43:391–415. doi:10.1146/annurev-neuro-100219-105424

McGinley MJ, David SV, McCormick DA. 2015. Cortical Membrane Potential Signature of Optimal States for Sensory Signal Detection. Neuron 87:179–192. doi:10.1016/j.neuron.2015.05.038

McIntyre CC, Grill WM. 2000. Selective microstimulation of central nervous system neurons. Ann Biomed Eng 28:219–233. doi:10.1114/1.262

Noei S, Zouridis IS, Logothetis NK, Panzeri S, Totah NK. 2020. Noradrenergic locus coeruleus ensembles evoke different states in rat prefrontal cortex. bioRxiv. doi:10.1101/2020.03.30.015354

Nowak LG, Bullier J. 1998. Axons, but not cell bodies, are activated by electrical stimulation in cortical gray matter. I. Exp Brain Res 118:489–500. doi:10.1007/s002210050305

O’Connor DH, Clack NG, Huber D, Komiyama T, Myers EW, Svoboda K. 2010. Vibrissa-based object localization in head-fixed mice. J Neurosci 30:1947–67. doi:10.1523/JNEUROSCI.3762-09.2010

O’Connor DH, Hires SA, Guo Z V, Li N, Yu J, Sun Q-Q, Huber D, Svoboda K. 2013. Neural coding during active somatosensation revealed using illusory touch. Nat Neurosci 16:958–65. doi:10.1038/nn.3419

Paxinos G, Watson C. 2007. The Rat Brain in Stereotaxic Coordinates. Academic, San Diego, USA.

Phillis JW, Kostopoulos GK. 1977. Activation of a noradrenergic pathway from the brain stem to rat cerebral cortex. Gen Pharmacol 8:207–11.

Poe GR, Foote S, Eschenko O, Johansen JP, Bouret S, Aston-Jones G, Harley CW, Manahan-Vaughan D, Weinshenker D, Valentino R, Berridge C, Chandler DJ, Waterhouse B, Sara SJ. 2020. Locus coeruleus: a new look at the blue spot. Nat Rev Neurosci 21:644–659. doi:10.1038/s41583-020-0360-9

Poulet JFA, Fernandez LMJ, Crochet S, Petersen CCH. 2012. Thalamic control of cortical states. Nat Neurosci 15:370–372. doi:10.1038/nn.3035

Ranjbar-Slamloo Y, Fazlali Z. 2020. Dopamine and Noradrenaline in the Brain; Overlapping or Dissociate Functions? Front Mol Neurosci 12:1–8. doi:10.3389/fnmol.2019.00334

Rodenkirch C, Liu Y, Schriver BJ, Wang Q. 2019. Locus coeruleus activation enhances thalamic feature selectivity via norepinephrine regulation of intrathalamic circuit dynamics. Nat Neurosci 22:120–133. doi:10.1038/s41593-018-0283-1

Sabri MM, Arabzadeh E. 2018. Information processing across behavioral states: Modes of operation and population dynamics in rodent sensory cortex. Neuroscience 368:214– 228. doi:10.1016/j.neuroscience.2017.09.016

Sachidhanandam S, Sreenivasan V, Kyriakatos A, Kremer Y, Petersen CCH. 2013. Membrane potential correlates of sensory perception in mouse barrel cortex. Nat Neurosci 16:1671–7. doi:10.1038/nn.3532

Safaai H, Neves R, Eschenko O, Logothetis NK, Panzeri S. 2015. Modeling the effect of locus coeruleus firing on cortical state dynamics and single-trial sensory processing. Proc Natl Acad Sci U S A 112:12834–12839. doi:10.1073/pnas.1516539112

Sara SJ. 2009. The locus coeruleus and noradrenergic modulation of cognition. Nat Rev Neurosci 10:211–23. doi:10.1038/nrn2573

Sara SJ, Bouret S. 2012. Orienting and reorienting: the locus coeruleus mediates cognition through arousal. Neuron 76:130–41. doi:10.1016/j.neuron.2012.09.011

Schriver BJ, Perkins SM, Sajda P, Wang Q. 2020. Interplay between components of pupil-linked phasic arousal and its role in driving behavioral choice in Go/No-Go perceptual decision-making. Psychophysiology 57:1–20. doi:10.1111/psyp.13565

Schwarz LA, Luo L. 2015. Organization of the Locus Coeruleus-Norepinephrine System. Curr Biol 25:R1051–R1056. doi:10.1016/j.cub.2015.09.039

Silvetti M, Lasaponara S, Horan M, Gottlieb J. 2021. A Reinforcement Meta-Learning Framework of Executive Function and Information Demand. bioRxiv 2021.07.18.452793. doi:10.1101/2021.07.18.452793

Silvetti M, Vassena E, Abrahamse E, Verguts T. 2017. Dorsal anterior cingulate-midbrain ensemble as a reinforcement meta-learner, bioRxiv. doi:10.1101/130195

Snow PJ, Andre P, Pompeiano O. 1999. Effects of locus coeruleus stimulation on the responses of SI neurons of the rat to controlled natural and electrical stimulation of the skin. Arch Ital Biol 137:1–28.

Stoney SDJ, Thompson WD, Asanuma H. 1968. Excitation of pyramidal tract cells by intracortical microstimulation: effective extent of stimulating current. J Neurophysiol 31:659–669.

Tehovnik EJ, Tolias a S, Sultan F, Slocum WM, Logothetis NK. 2006. Direct and indirect activation of cortical neurons by electrical microstimulation. J Neurophysiol 96:512– 521. doi:10.1152/jn.00126.2006

Totah NK, Neves RM, Panzeri S, Logothetis NK, Eschenko O. 2018. The Locus Coeruleus Is a Complex and Differentiated Neuromodulatory System. Neuron 99:1055–1068.e6. doi:10.1016/j.neuron.2018.07.037

Totah NKB, Logothetis NK, Eschenko O. 2019. Noradrenergic ensemble-based modulation of cognition over multiple timescales. Brain Res 1709:50–66. doi:10.1016/j.brainres.2018.12.031

Unakafova VA, Gail A. 2019. Comparing Open-Source Toolboxes for Processing and Analysis of Spike and Local Field Potentials Data. Front Neuroinform 13. doi:10.3389/fninf.2019.00057

van Vugt MK, Sederberg PB, Kahana MJ. 2007. Comparison of spectral analysis methods for characterizing brain oscillations. J Neurosci Methods 162:49–63. doi:10.1016/j.jneumeth.2006.12.004

Vazey EM, Moorman DE, Aston-Jones G. 2018. Phasic locus coeruleus activity regulates cortical encoding of salience information. Proc Natl Acad Sci U S A 115:E9439–E9448. doi:10.1073/pnas.1803716115

von Heimendahl M, Itskov PM, Arabzadeh E, Diamond ME. 2007. Neuronal activity in rat barrel cortex underlying texture discrimination. PLoS Biol 5:e305. doi:10.1371/journal.pbio.0050305

Waterhouse BD, Moises HC, Woodward DJ. 1998. Phasic activation of the locus coeruleus enhances responses of primary sensory cortical neurons to peripheral receptive field stimulation. Brain Res 790:33–44. doi:10.1016/S0006-8993(98)00117-6

Waterhouse BD, Navarra RL. 2019. The locus coeruleus-norepinephrine system and sensory signal processing: A historical review and current perspectives. Brain Res 1709:1–15. doi:10.1016/j.brainres.2018.08.032

Weiss E, Kann M, Wang Q. 2023. Neuromodulation of Neural Oscillations in Health and Disease. Biology (Basel) 12. doi:10.3390/biology12030371

Yang M, Logothetis NK, Eschenko O. 2021. Phasic activation of the locus coeruleus attenuates the acoustic startle response by increasing cortical arousal. Sci Rep 11:1–14. doi:10.1038/s41598-020-80703-5

Zabeh E, Foley NC, Jacobs J, Gottlieb JP. 2022. Traveling waves in the monkey frontoparietal network predict recent reward memory. bioRxiv 2022.02.03.478583. doi:10.1101/2022.02.03.478583

Zagha E, McCormick D a. 2014. Neural control of brain state. Curr Opin Neurobiol 29C:178–186. doi:10.1016/j.conb.2014.09.010

